# A Minimal, Adaptive Binning Scheme for Weighted Ensemble Simulations

**DOI:** 10.1101/2020.11.05.369744

**Authors:** Paul A. Torrillo, Anthony T. Bogetti, Lillian T. Chong

## Abstract

A promising approach for simulating rare events with rigorous kinetics is the weighted ensemble path sampling strategy. One challenge of this strategy is the division of configurational space into bins for sampling. Here we present a minimal adaptive binning (MAB) scheme for the automated, adaptive placement of bins along a progress coordinate within the framework of the weighted ensemble strategy. Results reveal that the MAB binning scheme, despite its simplicity, is more efficient than a manual, fixed binning scheme in generating transitions over large free energy barriers, generating a diversity of pathways, estimating rate constants, and sampling conformations. The scheme is general and extensible to any rare-events sampling strategy that employs progress coordinates.

## Introduction

Path sampling strategies have been pivotal in enabling the simulation of pathways and kinetics for rare events such as protein un(binding),^1–6^ protein (un)folding,^7–10^ and membrane permeation.^11,12^ These strategies exploit the fact that the time required to cross a free energy barrier (t_b_) is much shorter than the dwell time in the preceding stable (or metastable) state (t_b_ << t_dwell_) during which the system is “waiting” for a lucky transition over the barrier.^13,14^ By focusing the computational power on the actual transitions between stable states rather than on the stable states themselves, path sampling strategies can be orders of magnitude more efficient than standard simulations in sampling the functional transitions of rare events without introducing any bias into the dynamics.^15^

A major challenge for path sampling strategies has been the division of configurational space for a rare-event process. The application of these strategies can therefore be greatly streamlined by schemes that automate the adaptive placement of bins along a chosen progress coordinate. Such adaptive binning schemes have included the use of Voronoi bins^16–18^ and a variance-reduction approach^19^ for the weighted ensemble strategy;^20,21^ interfaces have also been used as “bins” to improve flux through bottlenecks between (meta)stable states^22–24^ for non-equilibrium umbrella sampling^23^ and forward flux sampling.^25^

Here, we present a minimal adaptive binning (MAB) scheme within the framework of the weighted ensemble strategy. The scheme can be used with high-dimensional progress coordinates and exhibits the following features: (i) no prior test simulations or training sets are required as the scheme relies only on the positions of the trailing and leading trajectories along the progress coordinate at chosen fixed time intervals; (ii) fewer bins are required compared with a manual binning scheme due to earlier identification of bottlenecks along the progress coordinate; (iii) the maximum number of CPUs (or GPUs) required is easily estimated prior to running the simulation since a similar number of bins are occupied throughout the simulation; and (iv) the scheme is easily extensible to more sophisticated schemes for adaptive binning.

To demonstrate the power of the adaptive binning scheme, we applied the algorithm to simulations of the following processes, in order of increasing complexity: (i) transitions between states in a double-well toy potential, (ii) molecular association of the Na^+^ and Cl^−^ ions, and (iii) conformational transitions of an N-terminal peptide fragment of the p53 tumor suppressor.

## Theory

### The weighted ensemble strategy

The weighted ensemble (WE) strategy involves running many trajectories in parallel and applying a resampling procedure at fixed time intervals *τ* to populate empty bins in configurational space--typically along a progress coordinate.^20,21^ The resampling procedure involves replicating trajectories that advance towards a target state, enriching for success in reaching the target state via a “statistical ratcheting” effect; to save computing time, trajectories that have not made any progress may be terminated, depending on which bin they occupy. Importantly, the rigorous tracking of trajectory weights ensures that *no bias is introduced into the dynamics,* thereby enabling the calculation of non-equilibrium observables such as rate constants. Furthermore, since the trajectory weights are independent of the progress coordinate, the progress coordinate as well as bin positions can be adjusted “on-the-fly” during a WE simulation.^16^

WE simulations can be carried out under non-equilibrium steady-state or equilibrium conditions.^26^ Non-equilibrium steady-state trajectories that reach the target state are “recycled” by terminating the trajectories and starting a new trajectory from the initial state with the same statistical weight. Equilibrium trajectories are not recycled, which means that target states need not be strictly defined in advance of the simulation.

### The MAB Scheme

The minimal adaptive binning (MAB) scheme works by first placing a fixed number of evenly spaced bins between “boundary” trajectories: the trailing and leading trajectories along the progress coordinate at a given time. After laying down these initial bins, the scheme identifies a “bottleneck” trajectory in each uphill direction of interest that maximizes the following objective function *Z* at a given time:

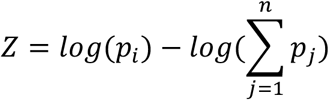

where *p*_*i*_ is the weight of trajectory *i* under consideration and 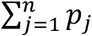 is the cumulative weight of all *n* trajectories that have surpassed trajectory *i* along the progress coordinate in the direction of interest. The scheme then assigns each boundary and bottleneck trajectory to a separate bin. In the final step, the WE strategy replicates and prunes trajectories within the same bin at fixed time intervals (WE iterations) to maintain a target number of trajectories per bin; trajectory weights are split or merged, respectively, according to rigorous statistical rules.^16^

**Figure 1** illustrates the steps of the MAB scheme for a one-dimensional progress coordinate:

1. Run dynamics for one WE iteration with a fixed interval *τ*.
2. Tag boundary and bottleneck trajectories with regard to the current WE iteration.
3. Adapt bin positions by dividing the progress coordinate evenly into a specified, fixed number of bins between the positions of the tagged trailing and leading trajectories; assign trailing, leading, and bottleneck trajectories to separate bins.
4. Replicate and prune trajectories to maintain a target number of trajectories in each bin.
5. Run dynamics with updated bins and repeat steps 1-4.

**Figure 1.**
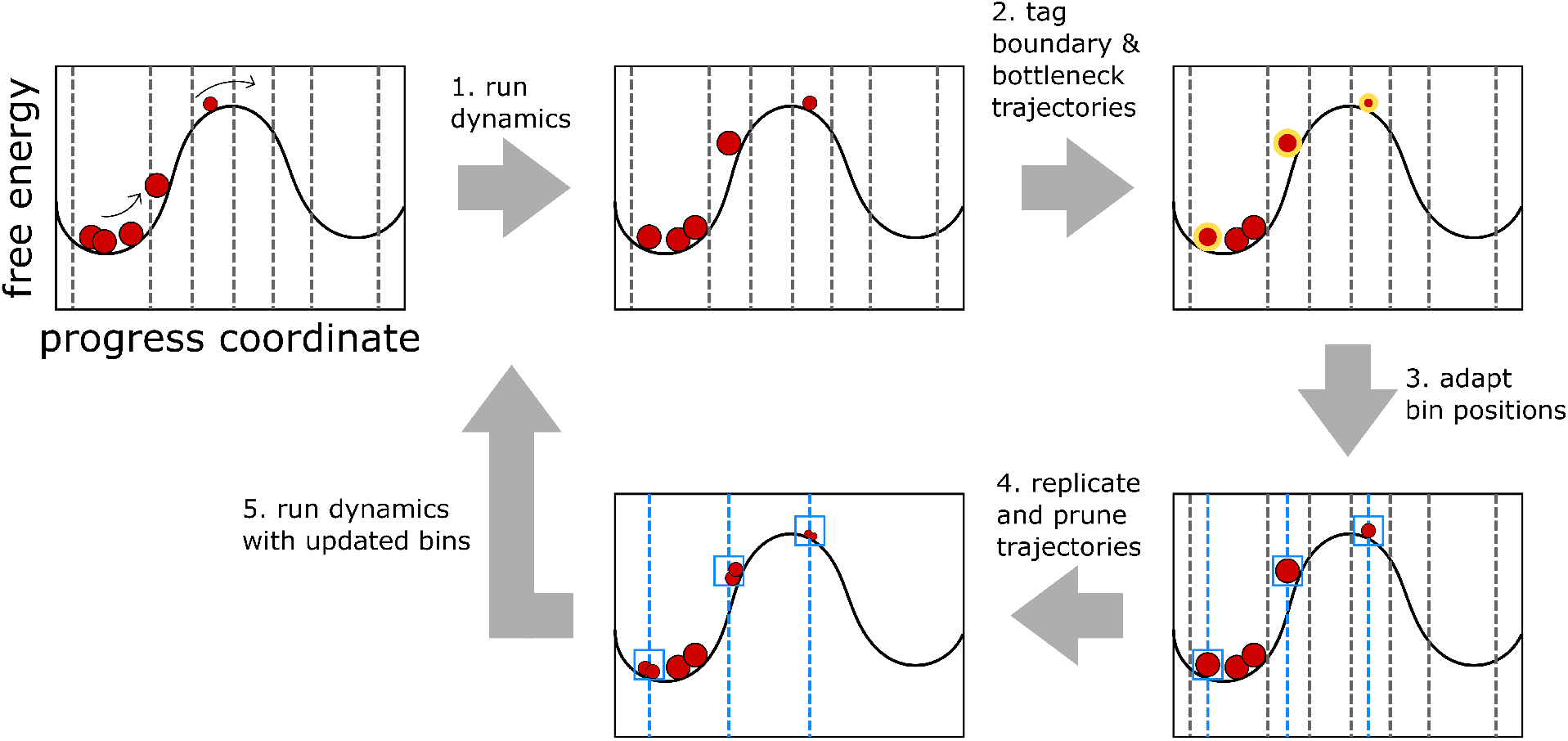
Illustration of the MAB Scheme for adaptive placement of bins along a one-dimensional progress coordinate. The scheme involves five steps. (1) Run dynamics for a short, fixed time interval *τ* using initial bins indicated by gray vertical lines. Trajectories are represented by red circles with sizes that are proportional to their statistical weights. (2) Tag boundary and bottleneck trajectories (highlighted in gold). (3) Adapt bin boundaries (blue vertical lines) by placing a fixed number of bins evenly between the positions of the trailing and leading trajectories along the progress coordinate and assigning each boundary and bottleneck trajectory to a separate bin (blue boxes). (4) Replicate and prune trajectories to maintain a target number of trajectories per bin. (5) Repeat steps 1-4 with updated bin positions until a desired amount of sampling is achieved.

For a multi-dimensional progress coordinate, steps 2 and 3 are carried out for each dimension of the progress coordinate. When multiple bottlenecks exist, replication of the most major bottleneck trajectory at the current WE iteration enriches for successful transitions over the corresponding bottleneck, enabling later bottlenecks along the landscape to be tackled. To avoid the replication of trajectories outside of the desired configurational space (*e.g.,* regions of unintentional protein unfolding), the MAB scheme includes the option to specify minimum and/or maximum limits of another observable as an additional dimension to the progress coordinate for the replication of trajectories. Since the number of trajectories per bin is fixed and a similar number of bins are occupied throughout the WE simulation—including separate bins for boundary and bottleneck trajectories—we can easily estimate the maximum number of CPUs (or GPUs) required for the simulation. A Python implementation of the MAB scheme is available for use with the WESTPA software^27^ (https://github.com/westpa/user_submitted_scripts/tree/master/Adaptive_Binning).

## Methods

### WE simulations

All WE simulations were carried out using the open-source WESTPA software package.^27^ For each benchmark system, we compared the efficiency of the MAB scheme for adaptive binning to a manual, fixed binning scheme. We present progress coordinates and binning schemes for each benchmark system below.

#### Double-well toy potential

The double-well toy potential consists of two equally stable states separated by a 34-kT free energy barrier. The potential was defined as

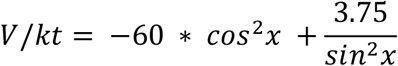

For the manual binning scheme, a one-dimensional progress coordinate was divided into 20 bins along a theoretical X position metric ranging from an initial state A at X=0.5 to a target state B at X=2.5; for the MAB scheme, a fixed number of bins ranging from 5 to 20 was used throughout the WE simulation for the same progress coordinate at any given time to determine the impact of the number of bins on the efficiency of generating successful transitions. For each binning scheme, a single WE simulation was run with a fixed time interval *τ* of 5 × 10^−5^ for each iteration and a target number of 5 trajectories/bin, yielding a total simulation time of 200,000 *δ*t.

Dynamics were propagated according to the overdamped Langevin equation:

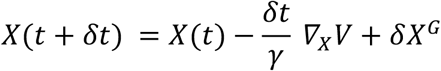

where *γ* is the friction coefficient, *δt* is the time step, and *δX*^*G*^ is a random displacement with zero mean and variance 2*γkTδt* with *δt* = 5 × 10^−5^ and reduced units of *γ* = 1and *kT* = 1.

#### Na^+^/Cl^−^ system

To sample Na^+^/Cl^−^ associations in explicit solvent, 5 independent, non-equilibrium steady-state WE simulations were carried out for each of the two binning schemes. A one-dimensional progress coordinate was used which consisted of the Na^+^/Cl^−^ separation distance. A total of 28 bins were equally spaced from a maximum value of 20 Å down to a target state at 2.6 Å. For both binning schemes, 1000 WE iterations were run with a fixed time interval *τ* of 2 ps for each iteration and a target number of 4 trajectories/bin, yielding an aggregate simulation time of 0.2 *μ*s.

Dynamics were propagated using the AMBER18 software package^28^ with the TIP3P water model^29^ and corresponding Joung and Cheatham ion parameters.^30^ Simulations were started from an unassociated state with a 12-Å Na^+^/Cl^−^ separation and a truncated octahedral box of explicit water molecules that was sufficiently large to provide a minimum 12-Å clearance between the ions and box walls. The temperature and pressure were maintained at 298 K and 1 atm using the Langevin thermostat (collision frequency of 1 ps^−1^) and Monte Carlo barostat (with 100 fs between attempts to adjust the system volume), respectively. Non-bonded interactions were truncated at 10 Å and long-range electrostatics were treated using the particle mesh Ewald method.^31^

#### P53 peptide

To sample alternate conformations of the p53 peptide (residues 17-29), a single equilibrium WE simulation was run using each of the two binning schemes and a two-dimensional progress coordinate that consisted of (i) the heavy-atom RMSD of the peptide from its MDM2-bound, *α*-helical conformation, and (ii) the end-to-end distance of the peptide. For both binning schemes, the WE simulations were run using a fixed time interval *τ* of 50 ps for each iteration and a target number of 4 trajectories/bin. A total simulation time of 2.0 *μ*s was generated for each binning scheme (338 and 200 WE iterations for the MAB and manual binning schemes, respectively). The MAB scheme used a maximum of 44 bins while the manual binning scheme used a maximum of 294 bins that were evenly spaced between an RMSD of 0 and 20 Å and end-to-end distance of 0 to 26 Å. For the MAB scheme, no other limits were specified for the replication of trajectories.

Dynamics were propagated using the AMBER18 software package^28^ with the Amber ff14SBonlysc force field^32^ and a generalized Born implicit solvent model (GBneck2 and mbondi3 intrinsic radii).^33^ Simulations were started from an energy-minimized conformation of the peptide that was based on the crystal structure of the MDM2-p53 peptide complex (PDB code: 1YCR).^34^ The temperature was maintained at 298 K using the Langevin thermostat and a collision frequency of 80 ps^−1^ for water-like viscosity.

### Standard simulations

A total of 5 independent 1-*μ*s standard MD simulations were run for the Na^+^/Cl^−^ system and a single 2-*μ*s simulation was carried out for the p53 peptide. Details of dynamics propagation and starting structures for these simulations are the same as those described above for the WE simulations.

### Calculation of rate constants

The association rate constant *k*^*RED*^ for the Na^+^/Cl^−^ system was directly calculated from the WE simulation using the Rate Event Duration (RED) scheme:^35^

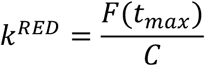

where the numerator *F*(*t*_*max*_) is the running average of the flux from the unassociated state to associated state over a steady-state WE simulation of total length *t*_*max*_; the denominator is a correction factor *C* that is equal to 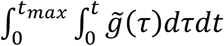, which incorporates the transient phase of the flux-evolution with time into the rate-constant estimation using the distribution 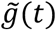 of event durations (barrier crossing times) during the interval of time [*t*, *t*_*max*_] in which it is possible to observe an event of duration time *t* from a simulation with a total length *t*_*max*_.

Uncertainties in the rate constants represent 95% confidence intervals, which is the standard error of the mean for each system multiplied by a critical value. For a large sample size (>30), this critical value would be 1.96, as obtained from a z-test. However, for the calculations in this study, which involve a smaller sample size (<30), critical values for determining the confidence interval at 95% were obtained from a t-test using the appropriate number of degrees of freedom (number of independent simulations minus 1) for each system.

### Estimation of WE efficiency in computing rate constants

The efficiency *S*_*k*_ of WE simulations in computing the association rate constant for the Na^+^/Cl^−^ system was estimated using the following:^20^

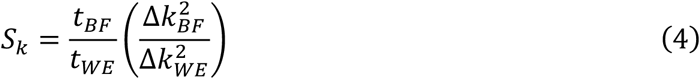

where *t*_*BF/WE*_ is the aggregate simulation time for standard “brute force” (BF) simulation or WE simulation, respectively, and Δ*k*_BF/WE_ is the relative error in the rate constants for the corresponding simulations where the absolute error is represented by the 95% confidence interval. Thus, the efficiency of the WE simulation in calculating the rate constant is determined by taking the ratio of aggregate times for the WE and brute force simulations that would be required to estimate the rate constant with the same relative error, with larger values of *S*_*k*_ corresponding to a more efficient simulation. The relative error in the rate constant is assumed to be inversely proportional to the simulation time.

## Results

We demonstrate the power of our minimal adaptive binning (MAB) scheme compared to fixed, manual binning schemes in weighted ensemble (WE) sampling of rare events. We applied the MAB scheme to the following processes, listed in order of increasing complexity: (i) transitions between stable states in a double-well toy potential, (ii) molecular associations of the Na^+^ and Cl^−^ ions, and (iii) conformational sampling of a peptide fragment of tumor suppressor p53.

### Simulations with a double-well toy potential

To test how effectively the MAB scheme performs for a process with a large free energy barrier, we focused on a double-well toy potential in which two equally stable states are separated by a 34 kT (20 kcal/mol at room temperature) barrier. WE simulations with a manual binning scheme (see **Fig. 2A** for bin positions) yielded no pathways from the initial state at *X* = 0.5 to the target state at *X* = 2.5 after 12,000 WE iterations, occupying only 14% of the fixed bins (**Fig. 2B**). In contrast, the MAB scheme generated pathways to the target state in 60 WE iterations, occupying 99% of the bins (**Fig. 2C**). This greater efficiency is due to the identification of bottleneck regions where the trajectory weights have sharply fallen. These regions correspond to the upward slope of the free energy barrier (**Fig. 2D**).

**Figure 2.**
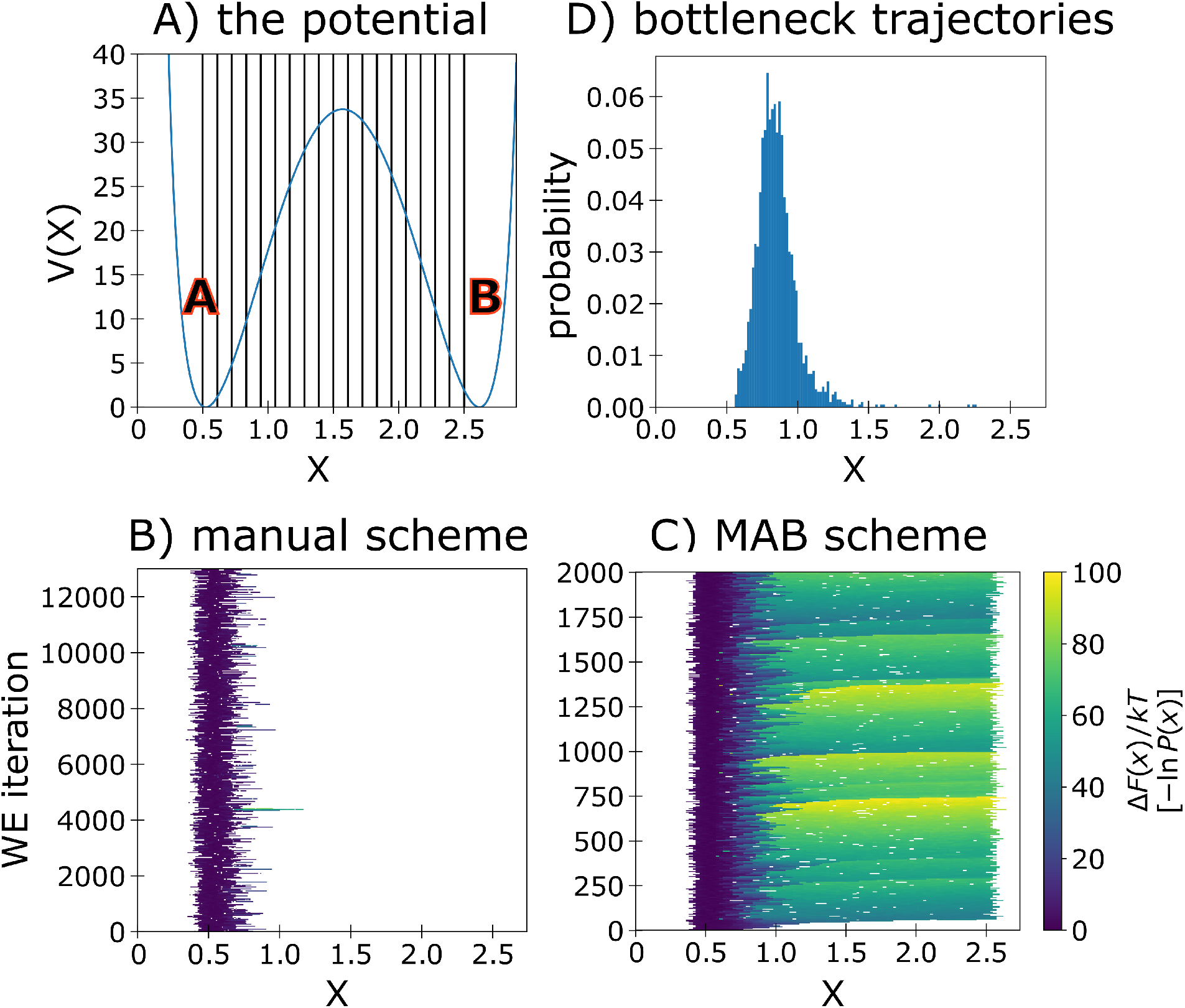
Transitions between stable states of a double-well toy potential. A) The double-well potential and manual binning scheme with 20 bins indicated by vertical lines. B) Probability distribution as a function of the WE iteration for a manual binning scheme. C) Probability distribution as a function of the WE iteration for the MAB scheme. D) Probability distribution of bottleneck walkers identified by the MAB scheme using 20 bins at any given time.

### Simulations of the Na^+^/Cl^−^ association process

To determine the effectiveness of the MAB scheme for a relatively fast process (ns timescale), we simulated the Na^+^/Cl^−^ association process in explicit solvent (**Fig. 3A**). Given the modest free energy barrier for this process, it was feasible to compute association rate constants using standard simulations, providing validation of the rate constants computed using WE simulations and manual/MAB binning schemes.

**Figure 3.**
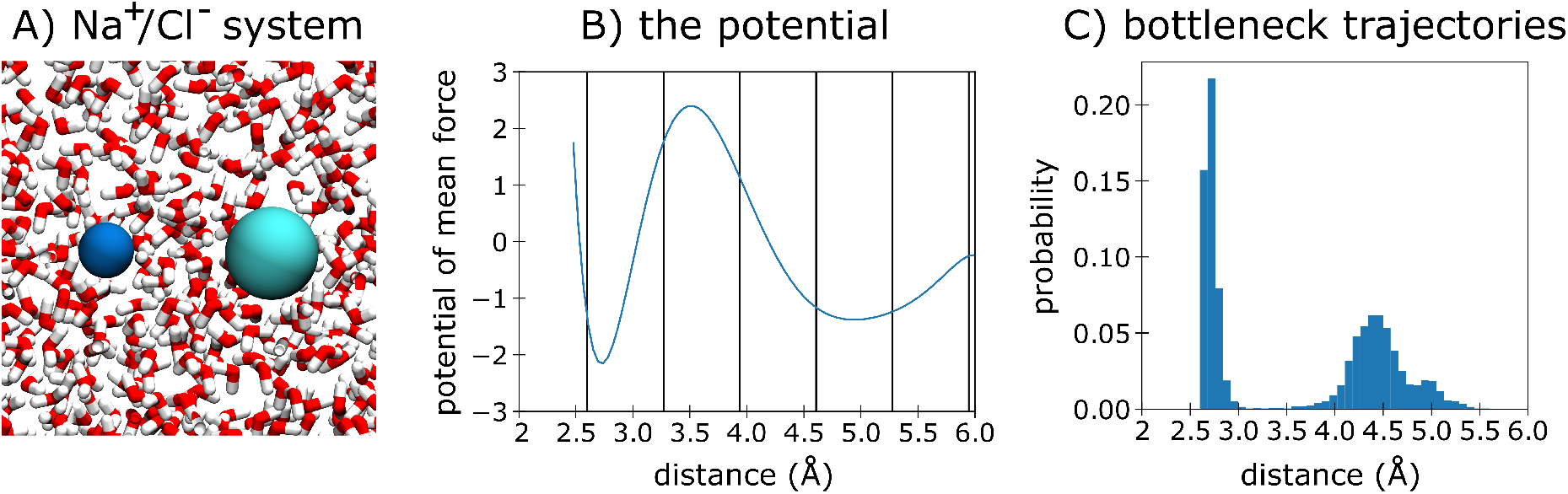
Molecular association of the Na^+^ and Cl^−^ ions. A) The Na^+^/Cl^−^ system in explicit solvent. B) Potential of mean force for the Na^+^/Cl^−^ association process with bin positions for the manual scheme indicated by vertical lines. C) Probability distribution of the positions of bottleneck trajectories tagged by the MAB scheme along the progress coordinate.

Table 1 shows the computed rate constants, efficiencies relative to standard simulations, and the number of successful pathways for WE simulations with the MAB and manual binning schemes. Regardless of the binning scheme, the WE simulations yield rate constants that are within error of the value from standard simulations (see also **Fig. S1**). Given the modest free energy barrier for this process, there is only a 1.6-fold gain in efficiency for the MAB scheme relative to the manual binning scheme (see **Fig. 3B** for bin positions). The MAB scheme also resulted in a 2-fold gain in the number of successful pathways related to the manual binning scheme.

**Table 1.** Computed rate constants for the Na^+^/Cl^−^ association process using WE simulations with the MAB scheme and manual binning scheme. Uncertainties represent 95% confidence intervals determined by a t-test. For each binning scheme, five WE simulations were run with each yielding 0.2 μs of total simulation time. The efficiency S_k_ of WE relative to standard simulations was calculated as described in Methods. For reference, the computed rate constant based on five 1-μs standard simulations was (3.9 ± 0.3) × 10^10^ M^−1^s^−1^.

| type of simulation | computed rate constant ($\text{M}^{-1}\text{s}^{-1}$) | total simulation time ($\mu\text{s}$) | $S_k$ | # successful pathways |
| --- | --- | --- | --- | --- |
| WE simulations with MAB scheme | $(3.9 \pm 0.3) \times 10^{10}$ | 1.0 | 5.1 | 2498 |
| WE simulations with manual binning scheme | $(4.1 \pm 0.4) \times 10^{10}$ | 1.0 | 3.1 | 1226 |

Consistent with our results for the double-well toy potential, the majority (60%) of the bottleneck trajectories occupied bins along the upward slope of the free energy barrier; the remaining bottleneck trajectories (40%) occupied bins located immediately before the target state (**Fig. 3C**).

### Conformational sampling of the p53 peptide

Given that the WE strategy has previously enhanced the conformational sampling of various biomolecules,^1,37^ we applied the MAB scheme to the conformational sampling of a p53 peptide (**Fig. 4A**). As expected, WE simulations using either the MAB scheme or the previously reported manual binning scheme^38^ yielded greater coverage of configurational space than standard simulations with the same total computing time (**Fig. 4B-C**). The MAB scheme placed bins more efficiently than the manual binning scheme, resulting in the occupation of 66% of the specified bins (29 out of 44 bins) compared to only 17% (50 out of 294 bins) for the manual binning scheme. Notably, the MAB scheme sampled a “horn shaped” region of the probability distribution which consists of primarily low-probability trajectories. This region was not sampled when using the manual binning scheme (or standard simulations) and includes a more extensive set of left-handed helices as well as PPII conformations, which have previously been identified as the dominant state by UV resonance Raman spectroscopy.^39^

**Figure 4.**
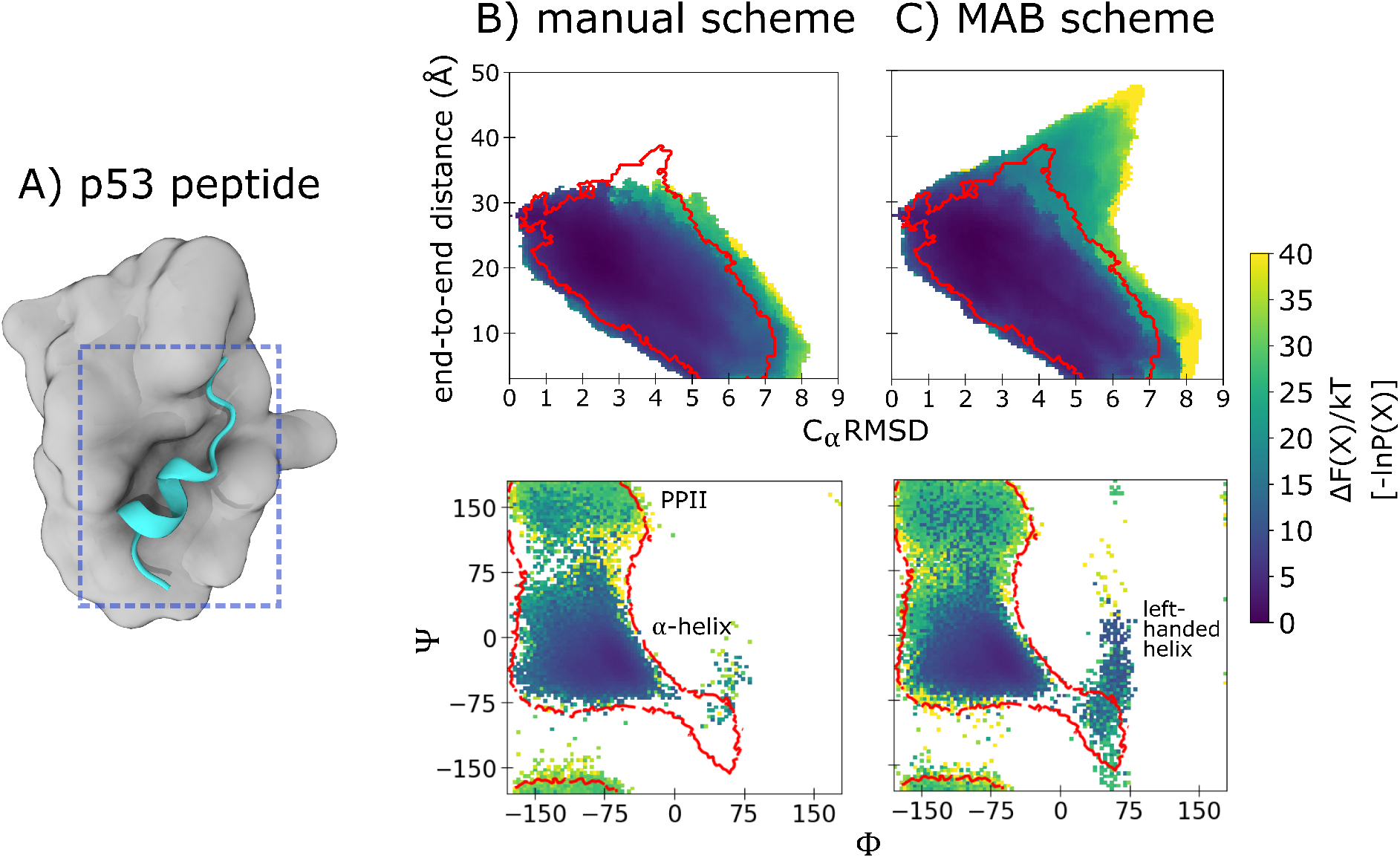
Conformational sampling of a p53 peptide. Probability distributions as a function of the two-dimensional WE progress coordinate from simulations of the p53 peptide (residues 19-23). **A)** starting conformation of the p53 peptide (cyan) for the WE simulation, extracted from the crystal structure^34^ of its complex with the MDM2 protein (gray). **B)** WE simulations using the manual binning scheme. **C)** WE simulations using the MAB scheme. Also shown are Ramachandran plots of the p53 peptide for the manual binning scheme and MAB scheme. Regions sampled by standard simulations are delineated in red.

## Discussion

On average, the minimal adaptive binning (MAB) scheme replicates more trajectories in steeper regions of the free energy landscape. As mentioned above for the double-well toy potential and Na^+^/Cl^−^ system (**Fig. 2D** and **Fig. 3C**, respectively), our MAB scheme identified bottleneck regions as the upward slopes of the free energy barriers, immediately before the barrier peaks (transition states). In contrast, a recently published variance-reduction strategy, which also seeks the most optimal placement of bins in weighted ensemble simulations, has identified such regions as the vicinity of the transition states, *i.e.* finer binning in transition-state regions yields the lowest variance in an observable of interest.^19^ This slight difference in the locations of the bottleneck regions is likely due to the fact that the goals of the MAB scheme and variance-reduction strategy are different. The MAB scheme aims to surmount free energy barriers whereas the variance-reduction strategy aims to minimize the variance of an observable of interest.^19^ Our results suggest that the MAB scheme would be particularly effective in surmounting large barriers when used with a “committor” coordinate,^13,40–43^ which tracks the probability that a given system configuration will commit to the target state before returning to the initial state—a nearly optimal, one-dimensional progress coordinate for the rare-event process of interest.^44,45^

The MAB scheme identifies bottleneck trajectories using an objective function that is easily extensible to track any arbitrary value. In its current form, the objective function tracks the probability of the trajectory in question along with the cumulative probability of all trajectories that are further along the progress coordinate of interest—all on a logarithmic scale. This requirement of having some trajectories that have surpassed the trajectory of interest makes it unlikely for identified bottleneck trajectories to be ones that have departed along orthogonal degrees of freedom (*i.e.* differentiating between a leading trajectory and a bottleneck trajectory). Alternatively, users may modify the objective function to track the average or maximum probability among trajectories that have surpassed the trajectory in question.

By identifying appropriate bin positions for use with other key WE parameters (*i.e.* resampling interval *τ* and target number of trajectories per bin), the MAB scheme greatly reduces the need for trial-and-error selection of these parameters, which are highly coupled to one another. To maximize the “statistical ratcheting” effect of the WE strategy, we recommend using the shortest possible *τ*-value that maintains high scaling of the WESTPA software with the number of GPUs (or CPU cores) on a given computing resource.^38^ Furthermore, our results indicate that a target number of either 4 or 5 trajectories per bin is sufficient to surmount large barriers or greatly enhance conformational sampling. If the goal is to simply generate pathways to a target state of interest, we recommend applying the MAB scheme with the minimal number of bins (*e.g.*, 5, with separate bins for the two boundary trajectories and one bottleneck trajectory in the direction of interest, and two bins between the boundary trajectories) to reduce the total computational time required for the simulation. Based on our tests with the double-well toy potential, the number of bins does not affect the ability of trajectories to reach the target state using the same total computing time (**Fig. S2**). However, if a greater diversity of trajectories or a rate-constant estimate is desired, we recommend applying the MAB scheme with a larger number of bins (15-20) to replicate more trajectories and yield more even coverage of configurational space along the progress coordinate.

## Conclusions

To streamline the execution of weighted ensemble (WE) simulations, we developed a minimal adaptive binning (MAB) scheme for automatically adjusting the positions of bins along a progress coordinate. Our scheme adjusts bin positions according to the positions of trailing, leading, and “bottleneck” trajectories at the current WE iteration. Despite its simplicity, the MAB scheme results in greater sampling of configurational space relative to manual binning schemes for all three benchmark processes of this study: (i) transitions between states of a double-well toy potential; (ii) Na^+^/Cl^−^ association; and (iii) conformational sampling of a peptide fragment of the tumor suppressor p53. Due to the earlier identification of bottlenecks along the progress coordinate, the MAB scheme enables the simulation of pathways for otherwise prohibitive large-barrier processes and a greater diversity of pathways when desired—all with dramatically fewer bins than manual binning schemes. As demonstrated previously, the efficiency of WE simulations relative to standard simulations is even greater for slower processes, increasing exponentially with the effective free energy barrier when the progress coordinate is appropriately binned.^46^

We recommend the MAB scheme as a general, minimal scheme for automating the placement of bins in combination with any rare-event sampling strategy that requires a progress coordinate. A particularly effective application of the scheme could be its use with a committor coordinate, which is a nearly optimal, one-dimensional progress coordinate for ordering states along simulated pathways for a process of interest according to a “kinetic ruler.” Regardless, the MAB scheme provides an ideal launching point for future developments of more sophisticated binning strategies by yielding initial, promising bins for further optimization.

## ASSOCIATED CONTENT

### Supporting Information

Figures S1 (time evolution of computed association rate constant for the Na^+^/Cl^−^ system) and S2 (probability distributions from WE simulations with the double-well potential and MAB scheme using different numbers of bins).

## Author Contributions

The manuscript was written through contributions of all authors. All authors have given approval to the final version of the manuscript.

## Notes

The authors declare the following competing financial interest: L.T.C. is an Open Science Fellow with Silicon Therapeutics.

## ACKNOWLEDGMENT

This work was supported by the NIH (1R01GM115805-01) and NSF (CHE-1807301) to L.T.C., and the University of Pittsburgh to P.A.T. (Honors College Brackenridge Undergraduate Research Fellowship and Dietrich School of Arts and Sciences Summer Undergraduate Research Award) and A.T.B (Arts and Sciences Graduate Fellowship). Computational resources were provided by the University of Pittsburgh’s Center for Research Computing. We thank Daniel Zuckerman (OHSU) for insightful discussions.

## ABBREVIATIONS

WE: weighted ensemble
MAB: minimal adaptive binning
BF: brute force

## TOC Figure

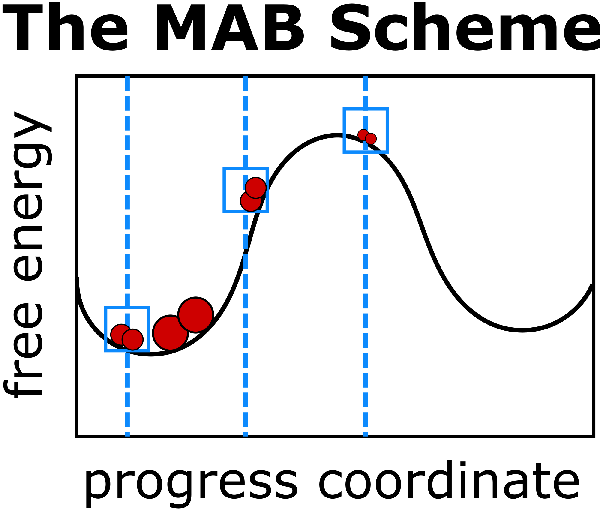

