## Supporting Information for "A Minimal, Adaptive Binning Scheme for Weighted Ensemble Simulations"

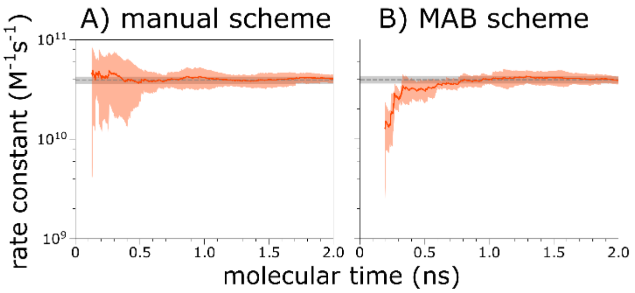


**Figure S1. Molecular association process of $Na^+$ and $Cl^-$ ions.** Computed rate constants for the molecular association process involving $Na^+$ and $Cl^-$ ions in explicit solvent as a function of molecular time $N\tau$ where $N$ is the number of WE iterations and $\tau$ is the fixed time interval for WE resampling. The rate constant from brute force simulations is shown with uncertainty as a shaded grey line. Results are shown for **A)** the manual binning scheme and **B)** the MAB scheme

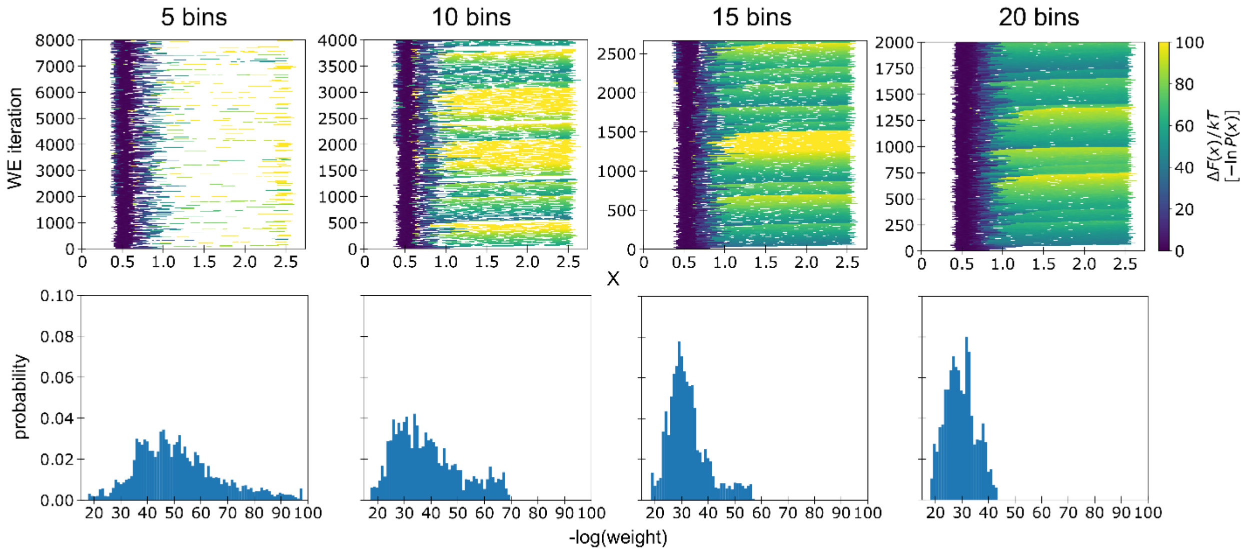


**Figure S2**. Probability distributions as a function of the progress coordinate X from WE simulations with the double-well potential and MAB scheme using different numbers of bins and the same total computing time (200,000 $\delta$t). Distributions of successful trajectories by their corresponding weights (trajectory weights shown on the logscale) are shown in the second row.
